# Models of increased latent HIV reservoir turnover prior to cART initiation imply novel clearance strategies

**DOI:** 10.1101/2021.06.06.447292

**Authors:** Aditya Jagarapu, Rajveer Mann, Michael J. Piovoso, Ryan Zurakowski

**Affiliations:** Department of Biomedical Engineering, University of Delaware, Newark, DE; Department of Electrical and Computer Engineering, University of Delaware, Newark, DE

## Abstract

CD4+ T cells with a naive or memory phenotype carrying a replication-competent HIV provirus are recognized as the major component of the persistent HIV reservoir. These cells only minimally express viral protein, reducing viral cytotoxicity effects and making them difficult targets for immune responses as well as every available antiretroviral drug. In patients on suppressive antiretroviral therapy, the half-life of these cells is approximately 4-5 years, balanced by clonal expansion of the cells resulting in an overall reservoir half-life in excess of 40 years. A recent study has shown that prior to the initiation of antiretroviral therapy, the half-life of these cells is instead on the order of two weeks. We present two models explaining the wide disparity in the on- and off-treatment half-lives of the quiescent infected T cells. In the first model, generalized (antigen non-specific) immune activation due to the high HIV viral loads explains the high latent reservoir turnover rates in the absence of treatment. If this mechanism dominates, we demonstrate that reduction of the latent reservoir size is possible, either through the administration of exogenous antigen or through the use of timed treatment interruptions. In the second model, direct killing of reservoir cells by HIV drives the increased turnover off-treatment. If this mechanism dominates, modulation of the reservoir size is not possible by the methods described above. Previously published models of the immune response to HIV show the possibility of inducing post-treatment control by reducing the latent reservoir size; by incorporating the same immune response dynamics in our first model, it is shown that it may be possible to induce post-treatment control using either exogenous antigen administration or timed treatment interruptions.

## 1 INTRODUCTION

Approximately, 27.4 million of the 37.6 million estimated persons living with HIV (PLWH) globally have access to combined antiretroviral therapy (cART) as of 2020. Modern cART effectively suppresses the viral replication in PLWH but is not curative. Lifelong treatment adherence is necessary to maintain a viral load below the detection limit, and any interruption to treatment leads to viral rebound. One of the major barriers in curing HIV is the persistence of long-lived naive and memory CD4+ T cells with integrated viral DNA that is replication competent but transcriptionally silent. Multiple factors contribute to transcriptional silencing in these cells, including the site of viral integration in the host DNA, epigenetic silencing, lack of cellular transcription factors, incomplete elongation of transcripts, nuclear retention of transcripts and micro RNAs limiting translation of viral proteins [7]. Existence of this repressive chromatin environment contributes to the stochastic reversion behavior of the latent reservoir cells, which randomly activate and begin producing virus, potentially re-seeding the viral pool even after years of suppressive therapy. Median estimates of the frequency of these latently infected cells is around 1 per million resting CD4+ T cells [8,11,19], but other studies have produced estimates ranging from as low as 0.03-3 [18] to as high as 55-108 [9] cells per million CD4+ T cells, depending on the type of measurement assay used and the requirement of virus functionality. Previous studies of the turnover of this reservoir, all of which followed successfully treated PLWH, observed slow turnover rates with an estimated half-life between 6 and 44 months, and an estimated time till total clearance around 70 years [5, 18].

Viral rebound due to the activation of these latent reservoir cells usually occurs within 2-3 weeks of treatment cessation. However, longer time to viral rebound was observed in some PLWH who received several years of cART early after infection [10, 17, 20]. PLWH who initiate cART early in primary infection preserve their innate immunity, maintain T and B cell diversity and function, exhibit accelerated immune responses, have limited viral diversity and exhibit reduced residual viral replication when compared to PLWH first treated during chronic infection. All these factors contribute towards extending the time to viral rebound, with values ranging from 1 – 8 years observed in the VISCONTI study [10]. A small subset of PLWH in these early treatment groups demonstrate a unique pattern. These PLWH, who exhibited high viral loads before initial treatment, maintain low viral loads (below detection limit) for an extended period following treatment cessation (possibly following an initial viral load spike), and are called post treatment controllers (PTCs). These PLWH have characteristics distinct from individuals termed as elite controllers (ECs) who possess strong immune responses and spontaneously maintain undetectable viral load without treatment [2,13,15]. Significant effort is going into developing treatment strategies that can alter the latent reservoir dynamics and induce post treatment control in PLWH. Approaches such as (i) “shock and kill”, where the HIV latency can be reversed through virus or immune mediated cytolysis and (ii) fully suppressing the HIV transcription leading to permanent silencing of virus production in the absence of cART is being extensively evaluated [6]. Latency revering agents (LRAs) are one such class of drugs that are being investigated that increase the rate of activation of these latent cells into infected cells thereby reversing latency [16] and producing virus which will be eventually targeted by ongoing cART therapy to avoid future replication cycles.

The existence of PTCs suggest that some PLWH might have two potential stable outcomes during off treatment conditions: one with a high viral load around 10^3^ – 10^5^ copies/ml corresponding to the standard outcome of PLWH and the other with a low viral load with measurements less than 50 copies/ml as observed in PTCs. Previous mathematical models developed by Conway et al., [4] considered the size of the latent reservoir, the rate of activation of the latent reservoir and the immune response as factors influencing the likelihood of a PTC outcome. Predictions from these models suggest that PTC can be achieved in PLWH having a small latent reservoir size and a high cell mediated immune response.

In order to determine when the latent reservoir is formed, Abrahams et al., [1] compared the sequences of replication competent viruses from resting CD4+ T cells on antiretroviral therapy to viral sequences circulating in the blood collected longitudinally prior to treatment initiation. They determined that a majority of the viruses from the latent reservoir were genetically similar to viruses replicating just prior to cART initiation. This experimental evidence suggests that the latent reservoir experiences continuous rapid turnover until treatment initiation and introduction of cART modifies the host environment in such a way that the latent HIV-infected cells stabilize. This evidence further leads us to hypothesize that the latent reservoir might have two distinct half-lives i.e. a shorter half life in the absence of treatment and a longer half life (44 months) during treatment conditions. A mechanism must exist by which the presence of high viral loads (i.e. during off treatment conditions) increases both the generation and turnover rates of the infected quiescent cell compartments. In this current study we introduce two possible mechanisms: 1) non-specific antigen driven activation of latent reservoir and 2) direct virus mediated reservoir killing. These two mechanisms are illustrated in Figure 1. We show that both mechanisms are capable of explaining the different turnover rates of the quiescent reservoir both on- and off-treatment, as is a combination of both mechanisms. Moreover, we show that if the increased turnover rate is driven by non-specific antigen driven activation, we show that modulation of the latent reservoir size is possible either through exogenous antigen infusion or timed treated interruptions. If the increased turnover rate is driven by direct virus mediated reservoir killing mechanism, we show modulation of the latent reservoir is not possible by these means. If a mix of above mechanisms drives the increased turnover rate, we predict reservoir modulation is possible depending on the relative contribution of each mechanism. Previous modeling work has tied the success of post-treatment control of HIV to the size of latent reservoir [4]. We demonstrate the possibility of designing treatment strategies to induce and stabilize post-treatment control, assuming dominance of mechanism 1.

**Fig. 1:**
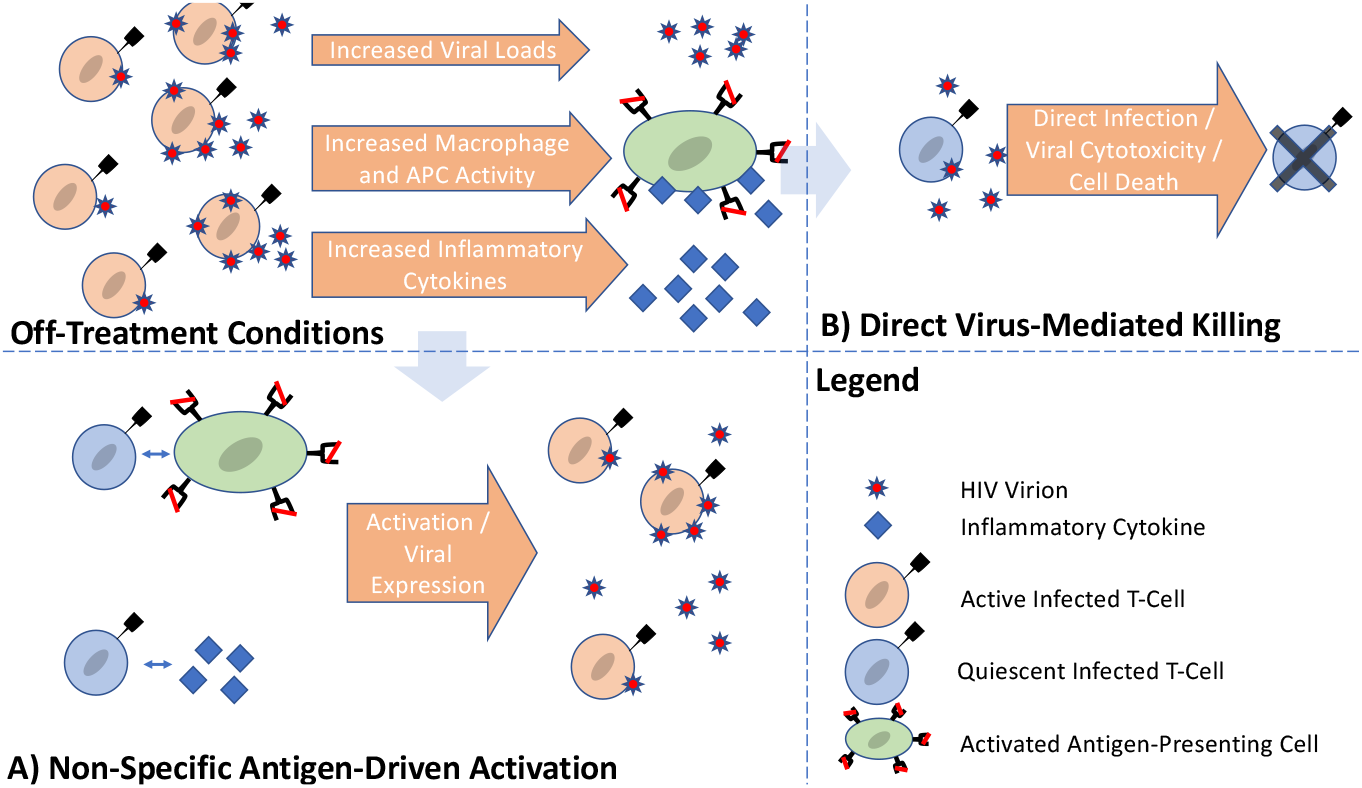
Off-treatment conditions lead to increases in viremia, activated macrophages, activated APCs, and inflammatory cytokines. Interactions with macrophages, APCs, and cytokines can increase turnover of quiescent infected cells by antigen non-specific activation (A). Direct interaction with the viremia can also increase turnover through direct virus-mediated killing due to infection and/or viral cytotoxicity (B).

## 2 MODELING RESERVOIR TURNOVER

The majority of the viral rebound after treatment interruption is due to the presence of latently infected CD4+ T cells. Understanding the formation, maintenance and turnover of this latent cell population is critical in order to design treatment strategies that can limit its growth and restrict viral rebound. The overall dynamics of the latent cell population (L) depends on the interaction between Uninfected/Target CD4 T cells (T), Virus (V) and the Infected CD4 T cells (I). These dynamics are captured in a previously published model [4] described in equations 1 to 4. Upon infection, a fraction (*α_L_*) of the target or uninfected CD4 T cells (T) become latently infected (L) and the remaining into infected cells (I). Target cells (T) are infected by the virus (V) at a rate proportional to their respective individual concentrations with density dependent infection rate constant ’*β*’ as shown in equation 1. Target cells are naturally formed at a constant rate ’*λ*’ and die at a rate ‘d’. Upon formation, the latent reservoir may proliferate at a rate ’*ρ*’, activate and produce infected CD4 T cells at a rate ’a’ and die at a rate ’*δ_L_*’. Productively infected cells (I) produce virus (V) at a rate ’p’ and the produced virus die at rate ’c’. The infected cells die at a rate ’*δ*’. The effect of antiretroviral therapy has been incorporated through drug efficacies, ’*ε_n_*’for drugs acting prior to nuclear integration, including Nucleoside, Non-Nucleoside and Integrase Inhibitors and ’*ε_p_*’ for drugs acting post nuclear integration, namely protease inhibitors. Protease inhibitors interrupt the maturation of viral particles, resulting in the formation of non-infectious pseudo-viral particles (P) that may nevertheless provide an antigenic stimulus. We assume these particles decay at a rate similar to that of free virus.

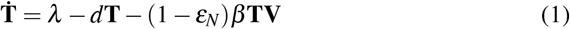

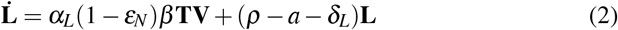

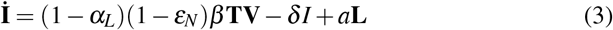

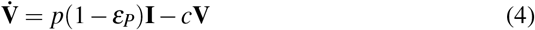

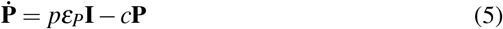

Figure 2 represents the compartmental model describing the interactions between various states of the above described dynamics.

**Fig. 2:**
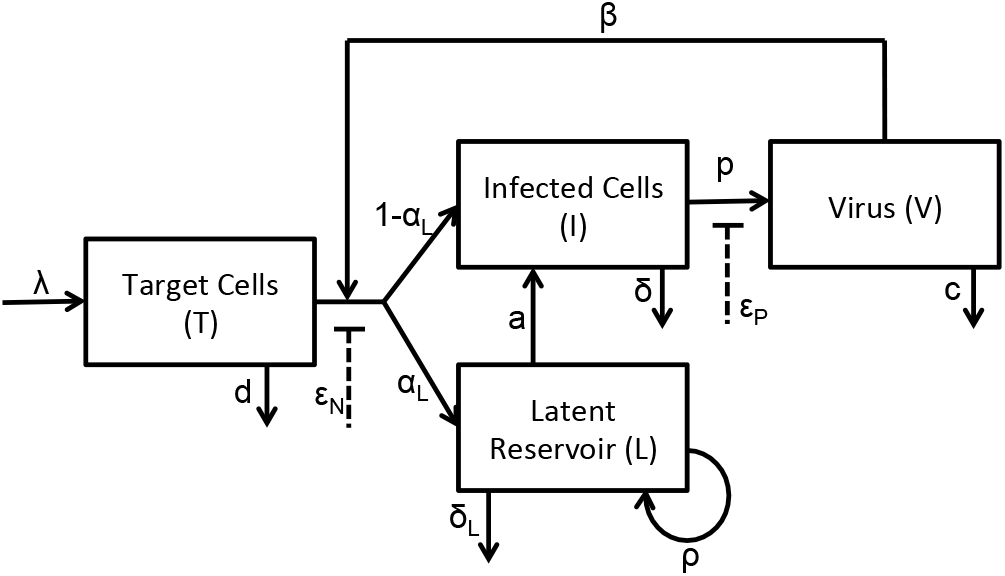
Mathematical model of HIV dynamics

Experimental studies on latent reservoir turnover by Abrahams et al. showed that the reservoir turnover prior to treatment initiation was orders of magnitude faster than the turnover rates observed during treatment [1]. The net growth rates of the reservoir, however, are very similar in both the on- and off-treatment conditions, implying that the mechanism increases both the creation and death rate of the reservoir in the absence of treatment. We propose two mechanisms that can explain the change in reservoir behavior by modifying the activation rates of latent cells in the previously published HIV models thereby incorporating the new experimental evidence. The first mechanism being discussed is the non-specific antigen-driven reservoir activation and the second mechanism is direct virus-mediated reservoir killing. Both the mechanisms are discussed as follows.

### 2.1 NON-SPECIFIC ANTIGEN-DRIVEN ACTIVATION

In this mechanism we assume that the presence of any non-specific antigen (i.e. viruses or virus mimetics) increases the rate of activation of the latent cells into infected cells. Previous studies have shown that pathogens (including virus and virus mimetics) bind to the receptors on the surface of antigen presenting cells (APCs) and activate a series of signals that result in the upregulation of costimulatory molecules (such as IL-15) on the surface of APCs that activate bystander T cells [3], [21].Since the viral load is high during off treatment conditions, we expect that the latent reservoir is under constant activation due to antigenic stimulation and the reservoir is constantly being regenerated through new infections, resulting in rapid proviral evolution. Once treatment is initiated, anti-retroviral therapy inhibits viral replication and the activation of latent reservoir is expected to reduce due to the lack of antigenic stimulation corresponding to low viral load. This decrease in the latent reservoir activation rate explains the low proviral evolution during on treatment conditions; consequently, the majority of the viral sequences sampled while on treatment are expected to match samples taken just prior to treatment initiation. As discussed above, the physiological mechanism involving antigenic stimulation involves activated APCs triggering bystander activation of CD4 T cells either directly or through the release of inflammatory cytokines; we model this behavior by incorporating Michaelis-Menten kinetics to the activation rate of the latent reservoir as shown in equations 7 and 8. ‘*K_max_*’ and ‘*K_h_*’ denote the latent reservoir maximum activation coefficient and the activation saturation coefficient respectively. ‘*K_min_*’ is considered as the basal activation rate in the absence of any antigen. The strength of the activation signal in this mechanism is expected to depend on the total antigen load, including both infectious virus *V* and non-infectious virus *P* as both are taken up by APCs.

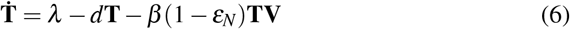

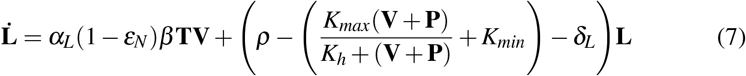

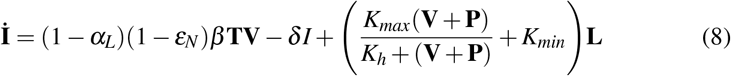

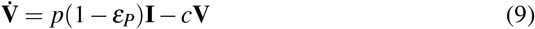

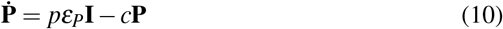

### 2.2 DIRECT VIRUS-MEDIATED RESERVOIR KILLING

In this mechanism, we assume that the difference in latent reservoir turnover during on and off treatment conditions is due to direct virus-mediated killing. Under this assumption, we hypothesize that the viral evolution during off treatment conditions is due to increased generation of latent cells due to new infections balanced by an increased death rate of these same cells. This increased death rate due to the virus-mediated killing can be modeled using second-order kinetics between the virus and the latent reservoir as shown in equation 12. In this mechanism, the pseudovirus (P) are irrelevant to the dynamics as they are non-infectious.

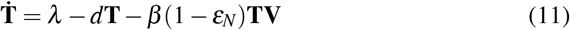

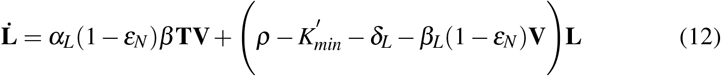

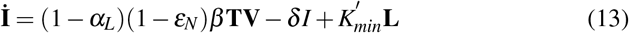

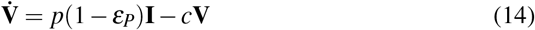

### 2.3 IMMUNE RESPONSE MODELING

In order to simulate the PTC phenomenon in the presence of pseudovirus and using treatment interruptions, we modified the previously published viral dynamics model proposed by Conway and Perelson [4]. Cytotoxic immune effector cells (E) are generated at a rate ’*λ_E_*’ and die at a rate ’*μ*’. Exposure to antigen from infected cells (I) stimulates clonal expansion of the effector cells, which proliferate at a Michaelis-Menten rate dependent on infected cell concentration (I) with ’*b_E_*’ as the maximum proliferation rate and ’*K_B_*’ as the 50% saturation coefficient. Immune cell exhaustion might also occur in the presence of high viral load. Immune cell exhaustion follows another Michaelis-Menten rate dependent on the infected cell concentration (I) with ’d*_E_*’ as the maximum exhaustion rate and *K_D_* as the 50% saturation coefficient for exhaustion. Effector cells (E) kill the productively infected cells (I) at a rate proportional to their respective concentrations with a proportionality constant ’m’ described as the effector cell killing rate. A high value of ’m’ corresponds to a strong immune response. Equations 15 through 20 describe the mathematical model (as shown in fig 3) corresponding to mechanism 1 under immune response settings. We also consider the possibility of infusing exogenous HIV antigen, similar to the naturally occurring pseudovirus (P), to promote the activation of latent reservoir during on treatment conditions. An input ’**u**’ is added to equation 19 to represent antigen infusion. Similar immune response modeling has been employed for the direct virus mediated killing mechanism. The equation for Effector cells (20) has been coupled to equations 11 through 14 in order to simulate mechanism 2 under immune response.

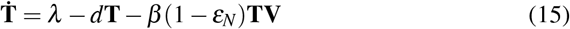

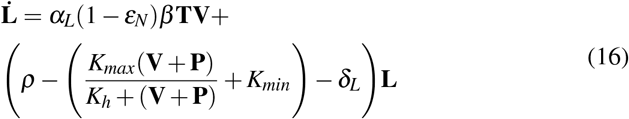

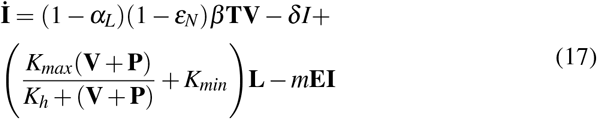

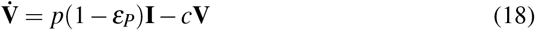

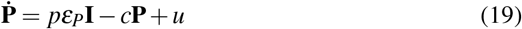

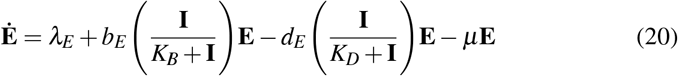

**Fig. 3:**
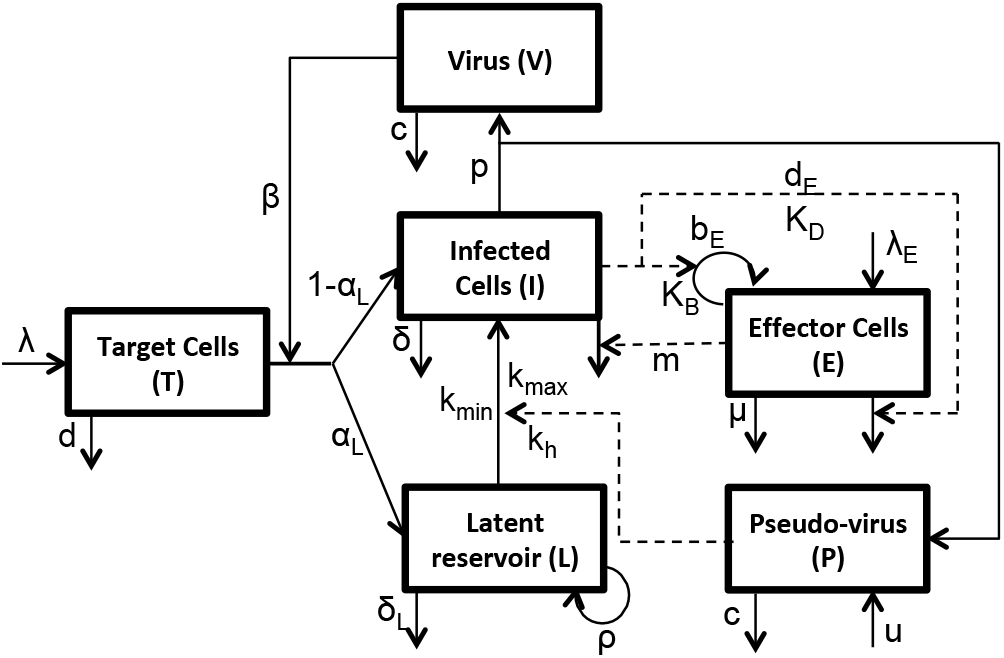
Mathematical model explaining virus dynamics and latency activation in the presence of immune response

**Fig. 4:**
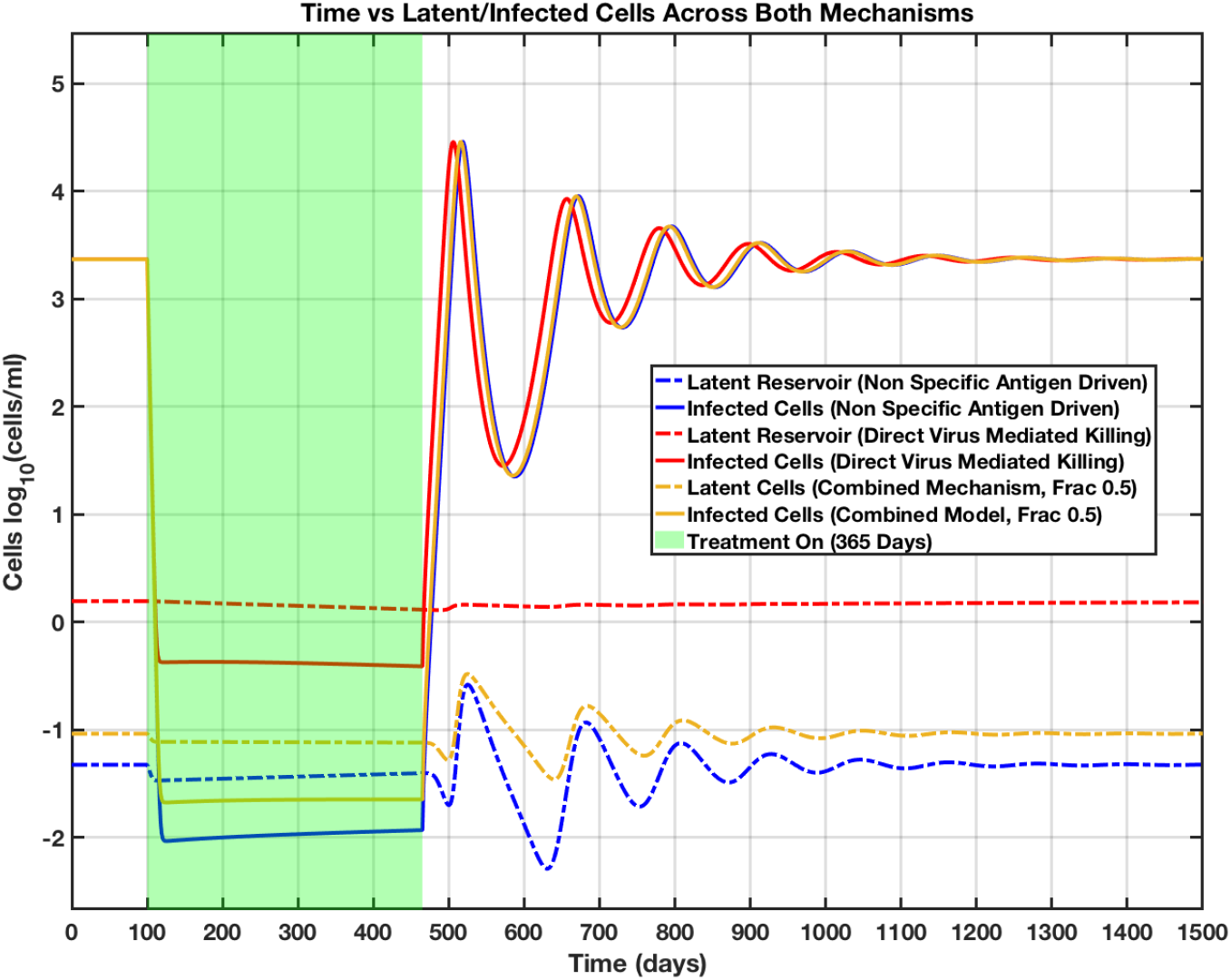
If Non-specific antigen-driven activation is present, the size of the latent reservoir oscillates following treatment interruption. Direct virus-mediated killing does not cause similar oscillations.

We have described two mechanisms, non-specific antigen-driven activation and direct virus-mediated killing, each of which independently explain the observed change in reservoir half-life when the patient is off treatment. It is also possible that both mechanisms might contribute towards the latent reservoir dynamics simultaneously in the same patient. This combined model incorporating both mechanisms can be described with the introduction of a new variable,’ *f*’ describing the relative contribution of each mechanism. The influence of this fraction *f* will regulate the dynamics for the latent and infected cells as shown in equations 21 and 22. The model equations for the remaining states (Target T cells, Virus, Pseudovirus and the Effector Cells) will be similar as discussed above in equations.

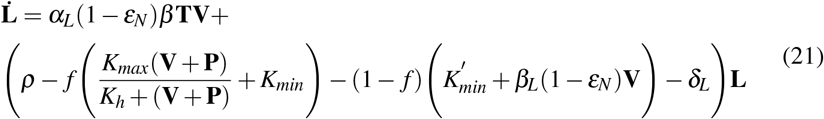

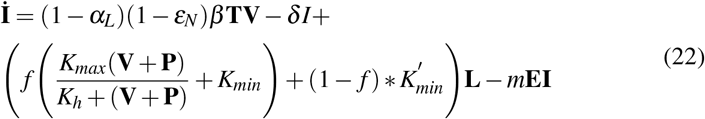

### 2.4 Parameters and Initial conditions

Parameters for the mathematical model including the basic HIV dynamics and the immune response are as previously reported [4]. Effector cell parameters corresponding to the PTC group with two stable steady states each corresponding to a high and low viral load have been identified and employed in our simulations. Baseline parameters for viral dynamics have been used as shown in table 1. Efficacy for the antiretroviral drugs have been assumed to be 0.5 for both NRTI and PI.

**Tab. 1:** VIRUS DYNAMICS MODEL PARAMETERS

| Definition | Units | Value |
| --- | --- | --- |
| Target cell production rate ($\lambda$) | cells/ml/day | $10^4$ |
| Target cells death rate ($d$) | 1/day | 0.01 |
| Density dependent infection rate ($\beta$) | ml/day | $1.5 \times 10^{-8}$ |
| Infected cell death rate ($\delta$) | 1/day | 1 |
| Virus production rate ($p$) | 1/day | 2000 |
| Virus clearance rate ($c$) | 1/day | 23 |
| Efficacy of NRTI ($\epsilon_N$) | - | 0.5 |
| Efficacy of PI ($\epsilon_P$) | - | 0.5 |

Parameters corresponding to the modified latent cell dynamics for both mechanisms (i.e. non-specific antigen driven activation and direct virus mediated reservoir killing) have been estimated based on the reservoir turnover rates during on and off treatment conditions observed by Abrahams et al. [1]. Previous studies assumed a constant value for the latent reservoir activation rate ’a’ at 0.001 *day*^−1^ [4]. In the modified latent cell dynamics, the same activation rate under non-specific antigen activation mechanism is assumed to follow hill function dynamics. The corresponding hill parameters *K_max_*, latent reservoir maximum activation coefficient and *K_h_*, latent reservoir activation saturation coefficient were obtained by solving the equation for latent reservoir half life corresponding to on/off treatment conditions as described below in equations 23 and 24. Similarly, for the direct virus mediated reservoir killing mechanism, modifications to the latent reservoir turnover rate has been implemented by introducing the parameter ’*β_L_*’, density dependent latent cell death rate as shown in equation 25 to simulate the enhanced death rate of the latent cells in the presence of virus. We used an estimated half life of 44 months for latent reservoir during on treatment conditions [18] and 2 weeks for off treatment conditions [1]. Substituting the half life values in equation 24 and 25 while assuming 10 and 10^7^ copies/ml as the viral load concentrations during on and off treatment conditions gave us a rough estimate for *K_max_* and *K_h_*, the hill function parameter values and *β_L_*, the density dependent infection rate for latent reservoir parameter. Estimates for *δ_L_* and *ρ* have been used from the same Conway and Perelson model [4]. In order to maintain a basal activation rate of the latent reservoir in the absence of virus (for both NSADA and DVMK mechanisms), a new parameter *K_min_* has been introduced. The value for *K_min_* has been adjusted depending on the mechanism under consideration. An updated estimate for *rho* at 0.2045 *day*^−1^ has been used to balance the net proliferation rate under both mechanisms. For the combined (NSADA and DVMK) mechanism, individual estimates for the corresponding parameters have been used. Estimated parameters for latent cell dynamics under non-antigen specific driven activation and direct virus mediated reservoir killing are shown in table 2 below.

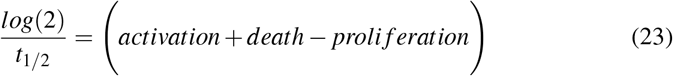

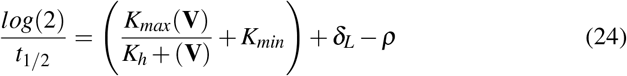

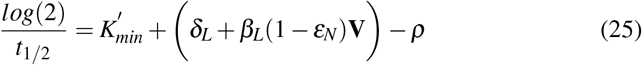

**Tab. 2:** LATENT RESERVOIR DYNAMICS PARAMETERS UNDER NSADA AND DVMK MECHANISMS

| *Definition* | *Units* | *Value* |
| --- | --- | --- |
| Latently infected cell death rate ($\delta_L$) | 1/day | 0.004 |
| Latently infected cell proliferation rate ($\rho$) | 1/day | 0.2045 |
| Fraction of newly infected cells that turn into latently infected cell ($\alpha_L$) | - | $10^{-6}$ |
| Latent reservoir maximum activation coefficient ($K_{max}$) [under NSADA] | 1/day | $5 \times 10^{-2}$ |
| Latent reservoir activation saturation coefficient ($K_h$) [under NSADA] | cells/day | 490 |
| Basal latent reservoir activation rate ($K_{min}$) [under NSADA] | 1/day | 0.2 |
| Density dependent latent cell death rate ($\beta_L$) [under DVMK] | ml/day | $4.9 * 10^{-9}$ |
| Basal latent reservoir activation rate ($K'_{min}$) [under DVMK] | 1/day | 0.201 |

Baseline parameters for immune response dynamics have been obtained from Conway and Perelson model [4]. Parameter regime corresponding to PTC predictions suggest that patients with intermediate effector cell response and low latent cell reservoir size were suitable candidates for PTC. Hence the effector cell killing rate ‘m’ has been chosen as 0.63 ml/cells/day in our simulations. Updated estimate for the effector cell proliferation coefficient *b_E_* at 0.1 *day*^−1^ has been used in order to obtain patient parameters with the possibility of a PTC behaviour for our modified latent cell activation model. Table 3 shows the patient parameters used for the immune response dynamics in our current simulations.

**Tab. 3:**
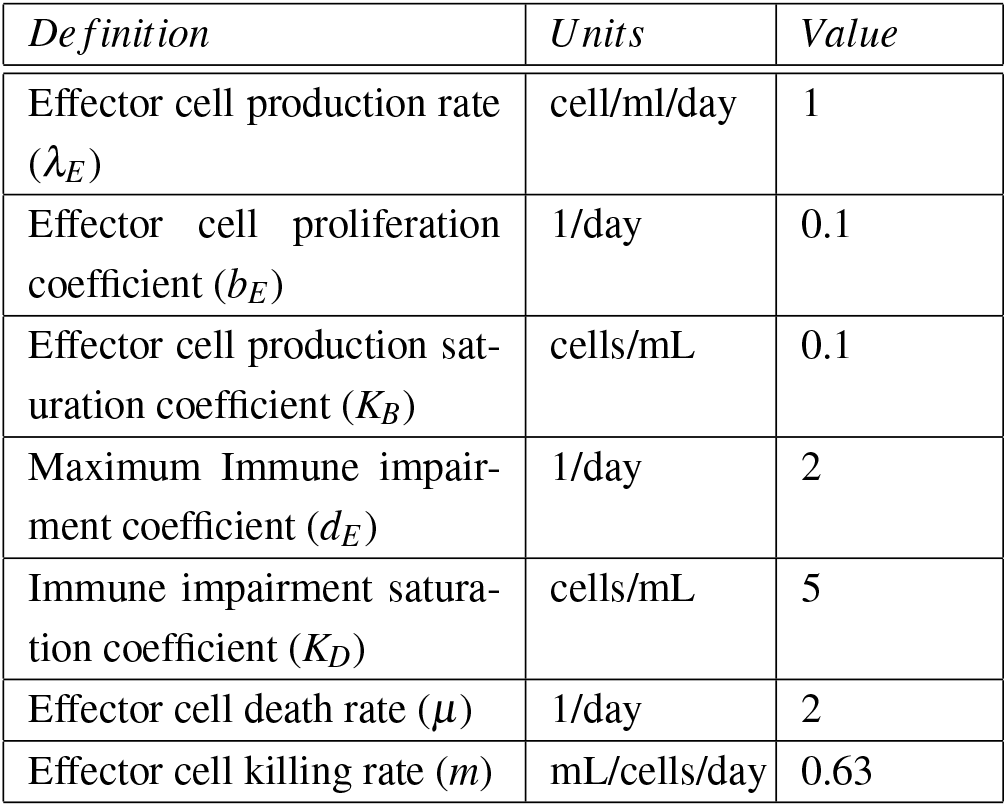
IMMUNE RESPONSE DYNAMICS PARAMETERS

Steady states corresponding to the above mentioned parameters have been obtained by solving model equations corresponding to the individual states depending on the mechanism under study. We used steady states corresponding to off-treatment conditions as initial conditions whenever the model has two steady states (i.e. one corresponding to high viral load and one belonging to low viral load).

## 3 RESULTS

### 3.1 BOTH MECHANISMS EXPLAIN VARIABLE RESERVOIR TURNOVER

Here we explain that non-specific antigen activation, direct virus-mediated killing, and any combination of the two are equally able to replicate the observed increase in reservoir turnover rate when the patient is off treatment.

### 3.2 MODULATION OF RESERVOIR SIZE BY TREATMENT INTERRUPTION OR ANTIGEN INFUSION

#### 3.2.1 Non-specific antigen-driven activation allows for modulation of reservoir size, but direct virus-mediated killing does not

Parameter values corresponding to HIV (table 1) and latent reservoir dynamics (Table 2) have been used to generate a virtual patient under both non-specific antigen driven (NSADA) and direct virus mediated killing (DVMK) mechanisms described in sections 2.1 and 2.2. The differences in the dynamics of latent reservoir between both mechanisms has been observed under on and off treatment conditions. Virtual patients in the current simulation have been subjected to HIV infection at day 0 and were maintained under off treatment conditions for an initial time period of 100 days. Combined antiretroviral therapy (cART) has been initiated to all the virtual patients at day 100 for a period of 1 year and were later withdrawn from cART indefinitely. Time evolution of latent reservoir and infected cells has been determined under on and off treatment conditions for a total time period of approximately 4 years as shown in figure 3. The latent reservoir dynamics corresponding to the patient under Non-Specific Antigen Driven Activation (NSADA) mechanism demonstrated conditions that would allow us to modulate the latent reservoir by introducing treatment at a later time point, specifically where the concentration of the latent reservoir is at its minimum value (in this current scenario that time point is around, t = 630th day). The oscillating behavior observed in the latent reservoir dynamics after treatment cessation under NSADA is due to the sudden activation of the latent reservoir due to the increase in virus concentration under off treatment producing more infected cells. The abrupt changes in the dynamics switching from on-treatment to off-treatment conditions under NSADA mechanism creates this oscillating behavior for the latent and infected cells (as seen in figure 3) which can be taken advantage of by switching off cART and initiating treatments at time points with the lowest concentrations of latent reservoir. However, the virtual patient illustrating Direct Virus Mediated Killing (DVMK) mechanism did not reveal any such oscillating dynamics that could be taken advantage of due to absence of antigenic stimulation. The absence of the activation component from the latent to the infected cell pool as a function of treatment conditions does not create any oscillatory behavior in the dynamics which could be taken advantage to lower the latent reservoir concentrations. Further, a majority of the patients in a population might not be having either of these mechanisms (i.e. NSADA and DVMK) acting independently. Hence, we considered a third virtual patient that could illustrate a combination of both these mechanisms according to equations 21 and 22 and evaluate the possibility of latent reservoir modulation under a combination of both these mechanisms. Depending on the extent of contribution by each mechanism, the modulation of latent reservoir is possible albeit to a less extent whenever DVMK dominates over NSADA mechanism in a particular patient. Figure 3 shows the behavior of all the three virtual patients illustrating conditions under which latent reservoir modulation can be achieved as described above under on and off treatment conditions.

#### 3.2.2 Treatment interruptions alone can achieve transient > 2 log_10_ reductions in reservoir size

The oscillatory behaviour observed in the latent pool under NSADA mechanism demonstrated the possibility of modulating the latent reservoir dynamics using treatment interruptions. We wanted to further investigate the extent to which we could reduce the latent reservoir concentrations using multiple treatment interruptions. As discussed above, we initiate treatment interruptions in virtual patients under pure NSADA and a combination of NSADA + DVMK with a fractional contribution of 50% from each mechanism, to estimate the lower bound of the latent reservoir pool that could be attained. Both the virtual patients were infected with HIV at day 0 and were kept under off treatment conditions for the initial 100 days. Anti-retroviral therapy was introduced at day 100 for a time period of 1 year and treatment interruptions were introduced after day 465. Treatment lengths spanning 1 day were introduced whenever treatment was initiated during the treatment interruptions phase. The treatment initiation schedule is determined using the following approach - for the first time point i.e. time at which we introduce treatment after day 465, we simulate the viral dynamics under the base treatment conditions (i.e. off treatment for the first 100 days, cART for the next 1 year and later off treatment for the rest of the time as discussed previously). We then estimate the time-point at which the latent reservoir achieves its lowest concentration under this base treatment schedule and introduce the next treatment (with length 1 day) at this time-point corresponding to this lowest concentration. Next, using this treatment schedule (i.e. base treatment schedule + first treatment initiation time-point), we simulate the viral dynamics again and introduce the second treatment at timepoint where the latent reservoir achieves its next lowest concentration value. We continue this iterative process until the latent reservoir saturates at its lowest achievable concentration with multiple treatment interruptions beyond which we cannot further lower the concentration by introducing more treatments. In this current scenario, we could modulate the concentration of latent reservoir as low as −3.73 log10 and −3.49 log10 concentrations under pure NSADA and combined NSADA + DVMK with fractional contribution of 50% from each mechanism. The treatment schedule consisted of 158 (for NSADA alone) and 450 (for combined NSADA+DVMK with fractional contribution of 0.5 for each mechanism) time-points at which treatment has been initiated in order to achieve this level of latent reservoir reduction under both the mechanisms. The fractional contribution from DVMK in the combined mechanism limits the latent reservoir from achieving similar levels of reservoir reduction when compared against a patient under pure NSADA mechanism. The latent reservoir reduction comparison between both patients under various degrees of mechanistic contributions can be seen in figure 5. The vertical lines at the bottom of each figure reflect the times at which the treatment is introduced in both the patients. The corresponding viral dynamics can also be seen in the lower plot under figure 5. We see that the viral levels rebound once cART has been permanently interrupted depicting observed cases of viral rebound during off treatment conditions.

**Fig. 5:**
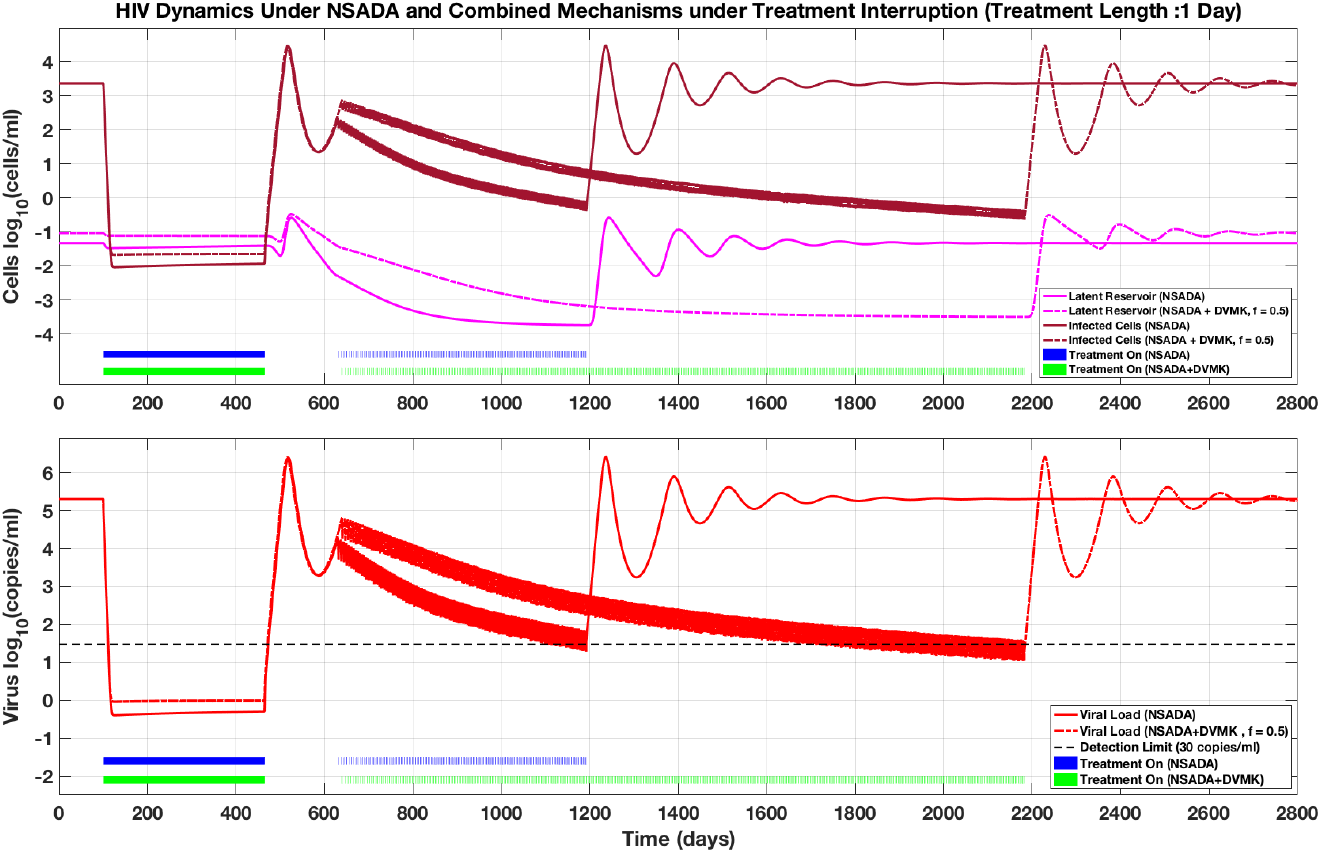
Treatment interruptions following discontinuation of continuous therapy leads to > 2 log reduction in reservoir size when NSADA dominates. Partial contribution of DVMK extends the time required to acheive reservoir reductions.

#### 3.2.3 Antigen infusion can achieve arbitrary reduction of reservoir size

In this current section, we introduce the concept of antigen infusion as an alternate strategy for modulating the latent reservoir dynamics. Banking on the hypothesis of antigen driven activation of the latent reservoir, we propose activating the latent reservoir using virus mimetics or pseudovirus (P) that are non-infectious that will aid in lowering the half-life of the latent reservoir during on-treatment conditions. This would allow the continuation of the antigen-driven activation of the latent reservoir pool in the presence of cART. Implementing the Hill function dynamics for the activation of latent reservoir in the presence of antigenic stimulation, we simulate the dynamics of the latent reservoir pool across various concentrations of pseudovirus (P) and varying levels of NSADA contribution in the combined mechanism as shown in figure 6. A total of nine combinations depicting nine virtual patients across three dosage values (1e3, 1e6 and 1e9 copies/ml) of pseudovirus and three levels of NSADA contribution (NSADA fractional contribution of 0.1, 0.5 and 0.9) in the combined mechanism have been used to understand the effect of pseudovirus influence on latent reservoir reduction. All the nine patients are assumed to be infected with HIV at day 0 and were kept under off-treatment conditions until day 100. Anti-retroviral therapy has been introduced at day 100 and were kept under on-treatment conditions until day 1000. Pseudovirus at various concentrations have been introduced between days 465 (i.e. approximately after one year of introducing cART) and 1000 along with cART. Simulations under these specified conditions reveal that the extent of latent reservoir reduction is highly influenced by the amount of dose and the extent of NSADA contribution in a particular patient. We observe that latent reservoir pool has been reduced by several orders of magnitude (−7 and −11.5 log_10_[per 10^6^ cells]) at higher doses of pseudovirus and also when the contribution of NSADA increased. However, in reality we might not observe latent reservoir concentrations less −3 log_10_[per 10^6^ cells] as stochastic processes dominate at lower concentration levels and the reservoir might get permanently silenced in some cases. We observe that the latent reservoir pool reach to original steady states after treatment interruptions depicting real case scenarios of viral rebound.

**Fig. 6:**
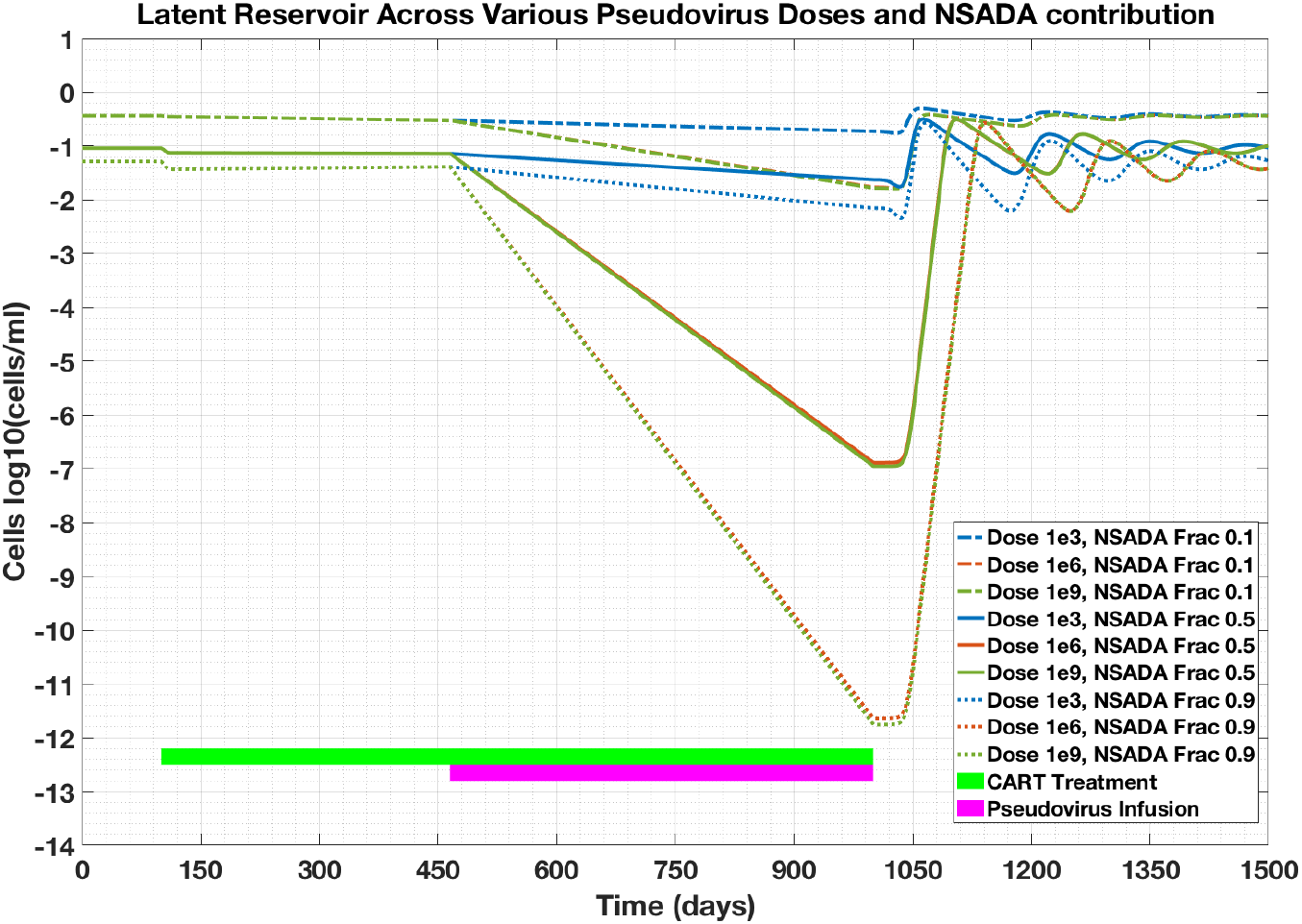
Infusing a non-infectious pseudovirus that provides antigenic stimulus could allow for reservoir reduction while on cART, with effect limited by dosage and duration of infusion.

#### 3.2.4 Increasing minimum treatmentment interruption time slows reservoir reduction

Previous treatment interruption simulations assumed a treatment length of 1 day in order to estimate the level of latent reservoir reduction under NSADA mechanism. In this current section, we evaluate conditions under which the treatment lengths vary between 1, 2, 7 and 14 days depicting the total number of days a patient would be on-treatment between treatment interruptions. Varying lengths of treatment has been assumed in order to evaluate the effect of cART adherence between treatment interruptions on modulating the size of latent reservoir. Figure 7 illustrates that the longer the treatment length the slower the rate at which the latent reservoir reduction is achieved. We also observe that with increase in treatment length, the saturation limit for the lower bound of latent reservoir reduction becomes higher.

**Fig. 7:**
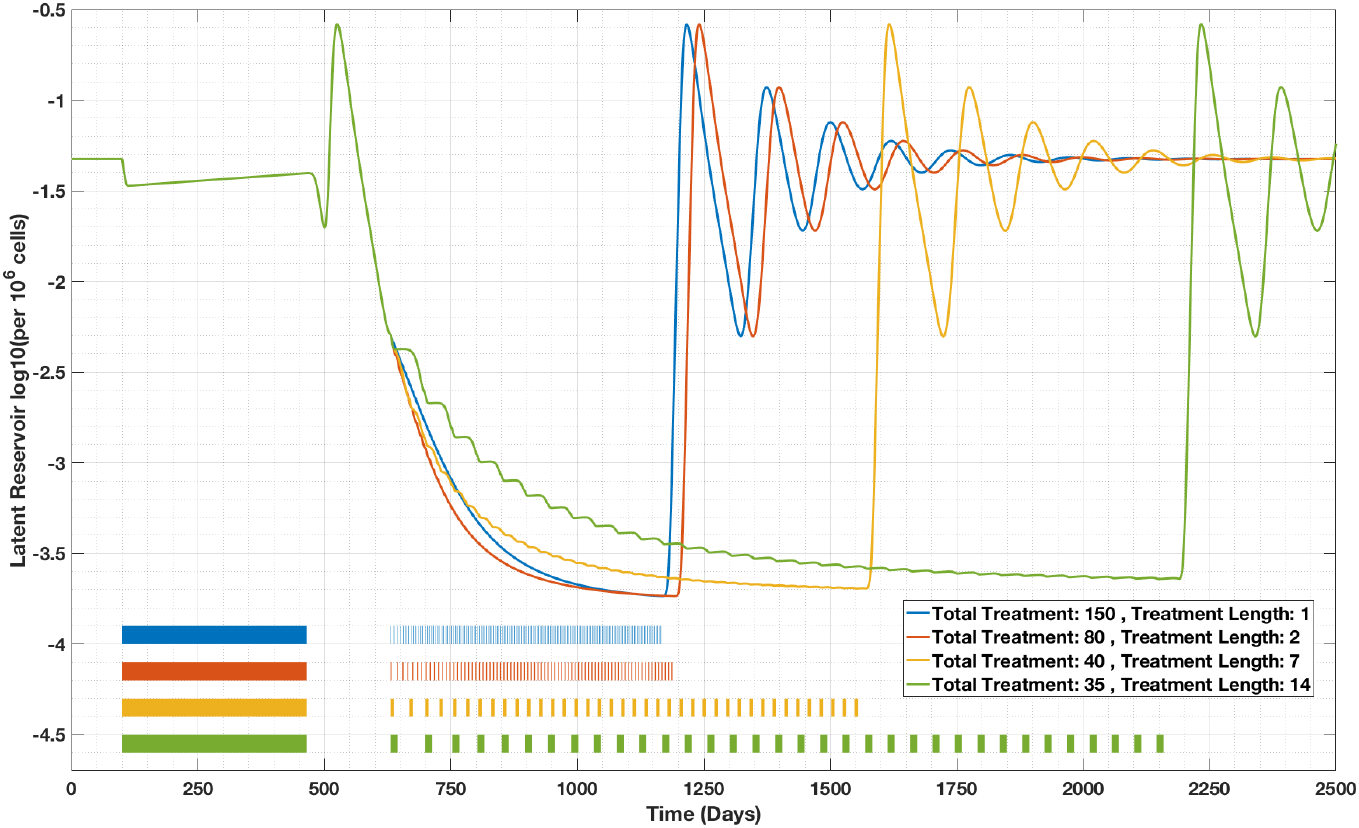
Imposing constraints on the frequency of treatment switching extends the time needed to achieve reservoir reduction.

#### 3.2.5 Constraints on maximum viral load slow reservoir reduction

Previous results on treatment interruption did not limit the viral rebound during treatment interruptions below a certain threshold. A consequence of this approach would result in conditions under which the virus might mutate and develop resistance against cART therapy eventually. In order to make sure that the virus does not escape the effects of anti-retroviral therapy, we introduced a constraint on viral rebound and evaluated the extent to which one could achieve similar levels of latent reservoir reduction as observed previously under non-constrained viral rebound conditions. Our simulations predict that similar levels of reservoir reduction is still possible. However, comparing the trajectories of the latent reservoir simulated under NSADA mechanism between figures 5 and 8 reveal that the total number of treatment interruptions required and the time it takes to achieve similar levels of latent reservoir reduction is more under constrained viral load conditions. A total of 1000 treatment interruptions were required to achieve −3.2 log_10_[per 10^6^ cells] reduction (under pure NSADA) in the latent reservoir pool over a time period of approximately 3285 days (i.e. approximately 9 years), while it took just 158 treatment interruptions over a time period of 735 days (i.e. approximately 2 years) in order to achieve −3.73 log_10_[per 10^6^ cells] reduction (under pure NSADA) under non-constrained viral load conditions.

**Fig. 8:**
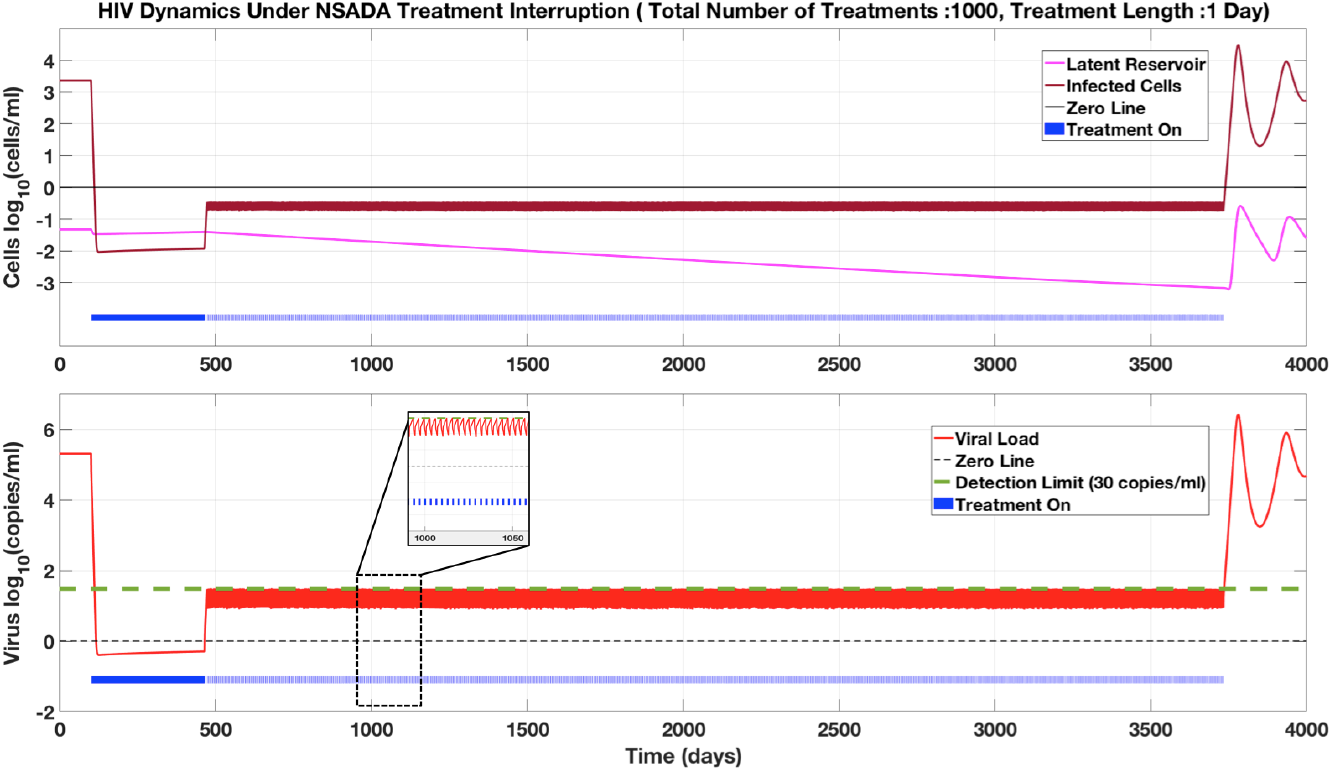
Imposing constraints on the maximum allowed viral load extends the time needed to achieve reservoir reduction.

### 3.3 STABILIZATION OF POST-TREATMENT CONTROL

In order to understand the effects of treatment interruption and antigen infusion in modulating the latent reservoir size and thereby influencing the formation of post treatment controllers, we simulate similar virtual patients as seen in section 3.2.2 and 3.2.3 in the presence of an immune response model. The mathematical equations governing the system of viral dynamics and the immune response model were previously described in section 2.3 under immune response modeling. The parameters for these simulations were obtained from tables 1, 2 and 3 under section 2.4 of parameters and initial conditions.

#### 3.3.1 Treatment interruptions following long-term suppressive therapy may stabilize immune control upon cART discontinuation

Three different scenarios were considered to understand the influence of the immune response on post treatment control behavior under treatment interruptions. Virtual patients under different latent reservoir turnover mechanisms i.e. pure NSADA and a combination mechanism with a fractional contribution of 0.5 from NSADA and DVMK were considered for this analysis as seen in figure 9. The total number of treatment interruptions were also varied between patients to understand its influence on the chances of a post treatment control. Initially, a virtual patient simulated under pure NSADA mechanism with 30 treatment interruptions did not show any post treatment controller behavior as the level of virus rebounded after the termination of treatment indefinitely. However, similar virtual patient i.e. patient with similar viral and immune model parameters exhibited post treatment controller behavior if the number of treatment interruptions were increased to 41 in number. We can see that the viral load corresponding to the virtual patient under 41 treatment interruptions (figure 9b) was below level of detection once the treatment was turned off indefinitely. Tracking the progression of immune response between these two patients help us answer the observed differences in the post treatment control behavior. Virtual patient under the higher number of treatment interruptions scenario could build the required immune strength over time (figure 9c) such that the developed immune response kept the viral load under control through cytolytic killing mechanisms. This critical immune strength could not be achieved in the patient with 30 treatment interruptions and hence there is a viral rebound after treatment interruption. In patients under combined turnover mechanism, we expect the total number of treatment interruptions to increase in number in order to achieve the post treatment control. Since the contribution of NSADA in the combined mechanism decreases, the activation of latent reservoir to infected cells decreases and hence it requires more effort to drive the latent reservoir to lower concentrations for the immune response to take charge and control the viral rebound. It required a total of 91 treatment interruptions for the virtual patient to show post treatment control behavior simulating combined latent reservoir turnover mechanism as seen in figure 8.

**Fig. 9:**
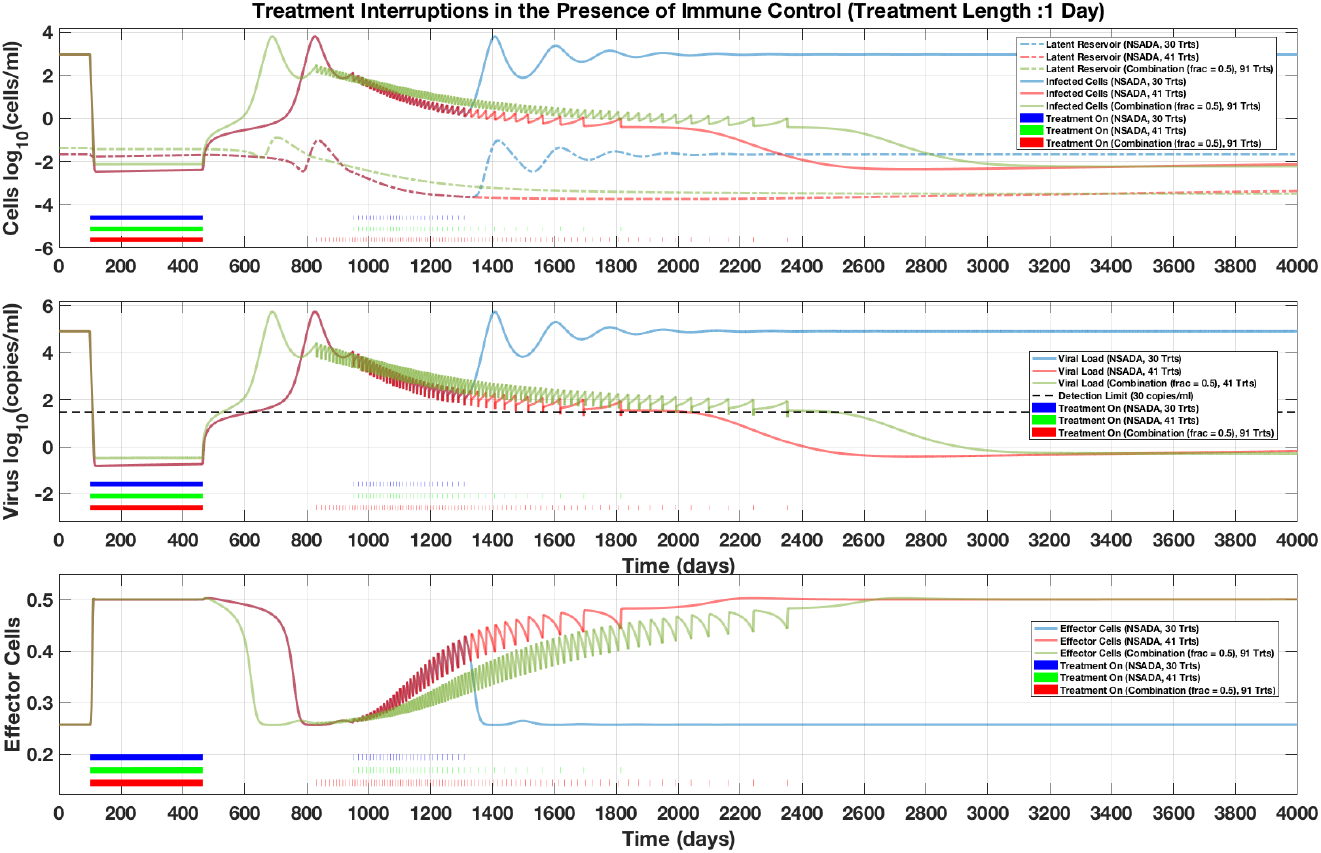
With immune response dynamics added to the model, well-chosen interruption schedules can stabilize post-treatment control of the virus (red, green), but not every interruption schedule leads to control (blue).

#### 3.3.2 Antigen infusion may stabilize immune control upon cART discontinuation

Similar analysis to treatment interruptions as discussed above was performed to virtual patients in the presence of antigen infusion. We wanted to understand the influence of antigen infusion on modulating the latent reservoir and the chances of observing post treatment control in the presence of immune response. Two different virtual patients were considered depending upon the level of NSADA contribution towards the latent reservoir turnover. One patient was simulated assuming pure NSADA mechanism and the other was simulated with a fractional contribution of 0.5 as shown in figure 10. Both the virtual patients were infused with pseudovirus at concentrations of 1e6 copies/ml for different treatment lengths. Both the virtual patients were subject to similar cART schedule (i.e. off treatment for the initial 100 days and then on-treatment for a period of 1 year) before the introduction of the cART+ pseudovirus treatment. For the individual under pure NSADA mechanism, a total of 4 days of antigen infusion was required in order to observe the post treatment control behavior. In the patient simulating under combined mechanism, a total of 39 days compared against 30 days of antigen infusion was required to observe the PTC phenomenon. The rational behind the difference observed between a PTC and a non-PTC behavior is whether the antigen infusion could equip itself to critical levels immune response or not in order to control the viral load from rebounding.

**Fig. 10:**
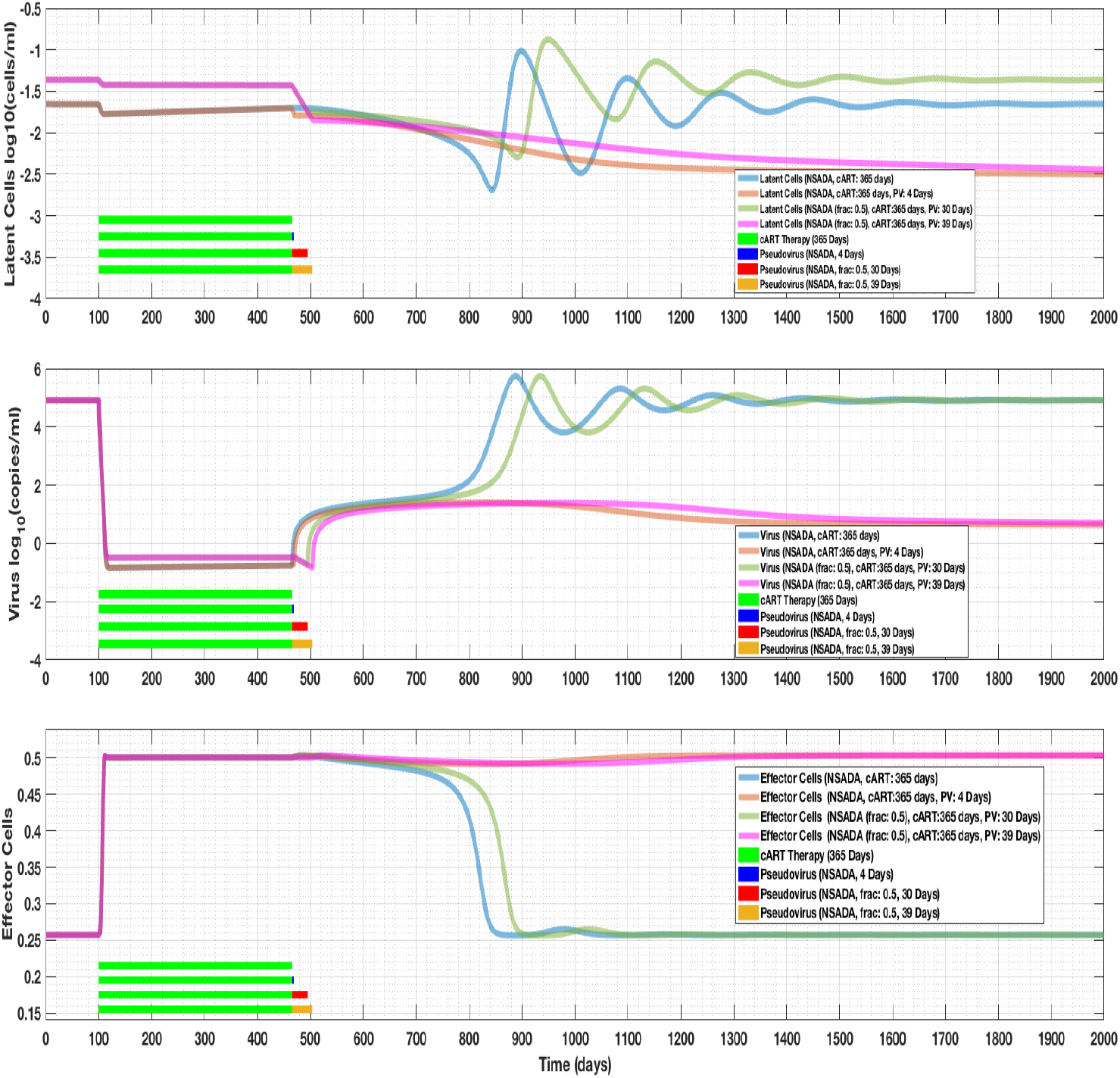
Short schedules of pseudovirus infusion can also result in stabilization of posttreatment control.

#### 3.3.3 Robustness analysis of open-loop post-treatment control stabilization schedules

In this section, we would like to demonstrate that the open-loop post treatment stabilization schedules is unique to patient and there is no single schedule for a population of patients. We demonstrate this idea by employing similar treatment interruption schedule for two virtual patients that have different levels of immune impairment during an immune response. For the patient with a lower immune impairment coefficient (i.e. *d_E_* = 2 1/d), a total of 41 treatment interruptions were required in order to achieve a post treatment control as shown in figure 11. However, similar treatment interruption schedule does not induce the post treatment control phenomenon in a patient with higher immune impairment coefficient value (i.e. *d_E_* = 3 1/d). A total of 64 treatment interruptions at different time-points were required in order to see the post treatment control behavior in the patient with higher immune impairment value. This demonstrates that each patient might be having a unique treatment interruption schedule that needs to be evaluated in order to achieve PTC.

**Fig. 11:**
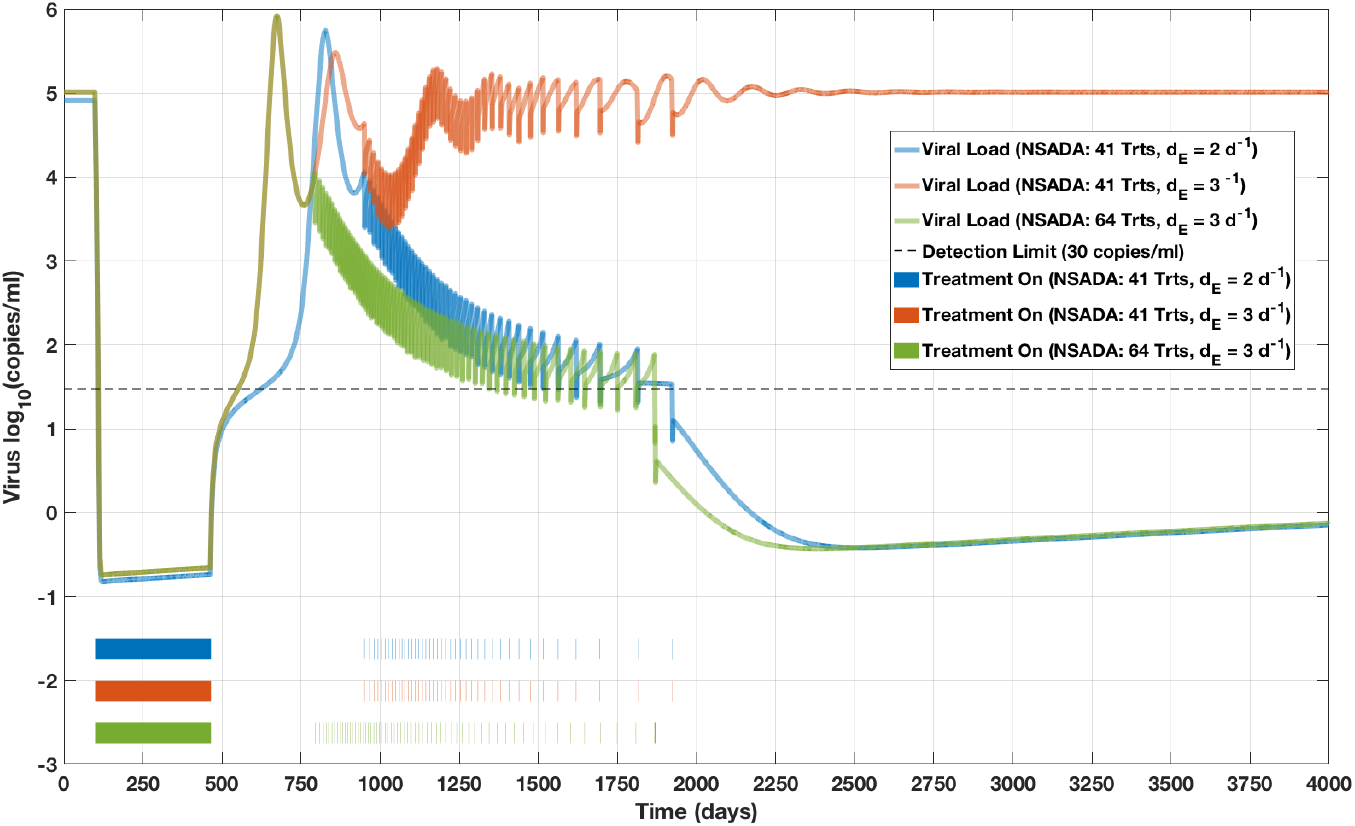
The treatment schedule that stabilized post-treatment control in one virtual patient (blue) does not stabilize PTC in another virtual patient with slightly different parameters (red), even though a treatment interruption schedule exists that will stabilize this second patient (green), illustrating the expected sensitivity of these methods to between-patient variations.

## 4 DISCUSSION

We have introduced a novel model of reservoir formation and turnover in the presence of antigens. This model is able to recapitulate the experimentally observed discrepancy between the long reported half-life of the quiescent HIV-infected reservoir during treatment and its apparent rapid turnover off treatment. This model furthermore introduces the possibility of modulating the turnover of the reservoir through the introduction of HIV viral antigen during treatment, which may be able to achieve substantial reductions in the size of the quiescent reservoir.

We coupled our model of reservoir dynamics with an existing model of the immune response to HIV, which had previously shown that post-treatment control depended on the size of the latent pool. The coupled model exhibited bistable behavior, with both post-treatment control and viral rebound as possible outcomes. Our models demonstrate that post-treatment control can be achieved in virtual patients with moderate immune response capability with the help of antigen infusion therapy. Optimization revealed that the shortest treatment period that can achieve post-treatment control for the parameter values used in our model is approximately 42 days, an extended but clinically achievable window. Future work should focus on clinical data-driven identification of the model parameters and development of robust strategies to facilitate immune-mediated post-treatment control. The coupling between quiescent cell turnover rate and virus-mediated immune activation should also be investigated in the context of recent evidence of ongoing HIV replication in pharmacological sanctuaries of treated patients [12, 14].

## ACKNOWLEDGMENTS

Research reported in this publication was supported by the National Institute Of Allergy And Infectious Diseases of the National Institutes of Health under Award Number R03AI136710. The content is solely the responsibility of the authors and does not necessarily represent the official views of the National Institutes of Health.

